# HyperMPNN – A general strategy to design thermostable proteins learned from hyperthermophiles

**DOI:** 10.1101/2024.11.26.625397

**Authors:** Moritz Ertelt, Phillip Schlegel, Max Beining, Leonard Kaysser, Jens Meiler, Clara T. Schoeder

## Abstract

Stability is a key factor to enable the use of recombinant proteins in therapeutic or biotechnological applications. Deep learning protein design approaches like ProteinMPNN have shown strong performance both in creating novel proteins or stabilizing existing ones. However, it is unlikely that the stability of the designs will significantly exceed that of the natural proteins in the training set, which are biophysically only marginally stable. Therefore, we collected predicted protein structures from hyperthermophiles, which differ substantially in their amino acid composition from mesophiles. Notably, ProteinMPNN fails to recover their unique amino acid composition. Here we show that a retrained network on predicted proteins from hyperthermophiles, termed HyperMPNN, not only recovers this unique amino acid composition but can also be applied to proteins from non-hyperthermophiles. Using this novel approach on a protein nanoparticle with a melting temperature of 65°C resulted in designs remaining stable at 95°C. In conclusion, we created a new way to design highly thermostable proteins through self-supervised learning on data from hyperthermophiles.

## Introduction

Proteins exist in nature in a delicate balance between function and stability, tolerating non-ideal mutations as long as function is maintained. In fact, some regions of the protein need to be thermodynamically frustrated on order to perform its function^1,2^. Therefore, natural proteins tend to be only marginally stable^3^. This is sufficient for natural proteins where there is no evolutionary benefit of the protein melting point greatly exceeding the optimal growth temperature (OGT) of its organism. In contrast, energetic frustration resulting in flexibility is a pre-requisite for protein function, as observed for catalytic residues in enyzmes^2^, and is tied to the physiological temperature of the source organism. However, industrial processes operate at temperatures exceeding OGTs of mesophile organisms to enhance reaction rates and reduce the risk of microbial contamination. The use of proteins as therapeutics or biocatalysts is therefore often limited by their initial stability and improving it is a typical goal of protein engineering campaigns.

Computational prediction of stabilizing mutations commonly makes use of either evolutionary data^4^, biophysical scoring functions^5,6^, or supervised learning^7,8^. Recent developments in high-throughput stability measurements^9^ have led to improved accuracy of deep learning approaches for the prediction of stabilizing mutations^7^. However, a caveat of this work is that strong generalization beyond the measured ‘mini-proteins with less than 100 amino acids, remains difficult. Additionally, improving stability of a protein beyond single digit degrees will require considerable redesign, while most available data refer to single-point mutations, not necessarily taking combination of mutations into account. On the experimental side, labor intensive rounds of directed evolution or mutagenesis studies can be used to stabilize a protein.

However, nature has already found general strategies for thermal stability in organisms adapted to extreme environments. Hyperthermophiles, living in thermal vents or hot springs, grow optimally at up to 105°C and their proteins have drastically higher average melting temperatures^10^. *Thermotoga thermophilus*, for example, has an OGT of around 70°C with an average protein melting temperature of 85°C which is 35°C higher than observed in *Escherichia coli (E. coli)*^11^. Multiple adaptations enable the high melting points, mainly an increase in hydrophobic core residues as well as charged surface residues while decreasing polar surface residues^10,11^. In contrast to enzymes from mesophiles, enzymes from hyperthermophiles have their catalytic optimum at high temperatures which are ideal for industrial processes, but tend to lose their catalytic properties at lower temperatures as a consequence of decreased flexibility required for function^12^.

Protein nanoparticles offer versatile promising biomedical applications, including drug delivery and vaccine development^13^. Thermostability is considered a crucial characteristic to extend their practical utility in real-world applications. The development of vaccines capable of withstanding high temperatures for extended periods is of great interest in the context of shelf-life and delivering vaccines in regions where uninterrupted cooling chains are challenging. Self-assembling protein nanoparticles (SAPN) are regarded as a promising platform due to their distinctive architecture. The I53-50 SAPN is characterized by its large lumen, multicomponent nature and controllable assembly, and has been successfully demonstrated in various vaccine candidates^14^.

In this work, we set out to establish a general strategy for the design of stable proteins inspired by hyperthermophiles. Therefore, we leveraged self-supervised learning by creating a dataset of predicted protein structures from hyperthermophiles which have a unique amino acid composition, and retraining ProteinMPNN^15^ to recover this composition. Next, we selected the I53-50 SAPN to experimentally verify the potential of HyperMPNN to design thermostable proteins.

## Results

### Collecting a dataset of predicted protein structures from hyperthermophilic organisms

Protein sequences were collected from a list of hyperthermophiles^16,17^ through UniProt^18^ (**Fig. 1**). All 96,738 sequences were clustered to 50% sequence identity resulting in 34,759 sequences. Subsequently, their predicted structures were downloaded from the AlphaFold2 database^19,20^. Filtering out structures with low AlphaFold model confidence (pLDDT cutoff < 70) resulted in 29,042 predicted structures of proteins from hyperthermophiles.

**Fig. 1:**
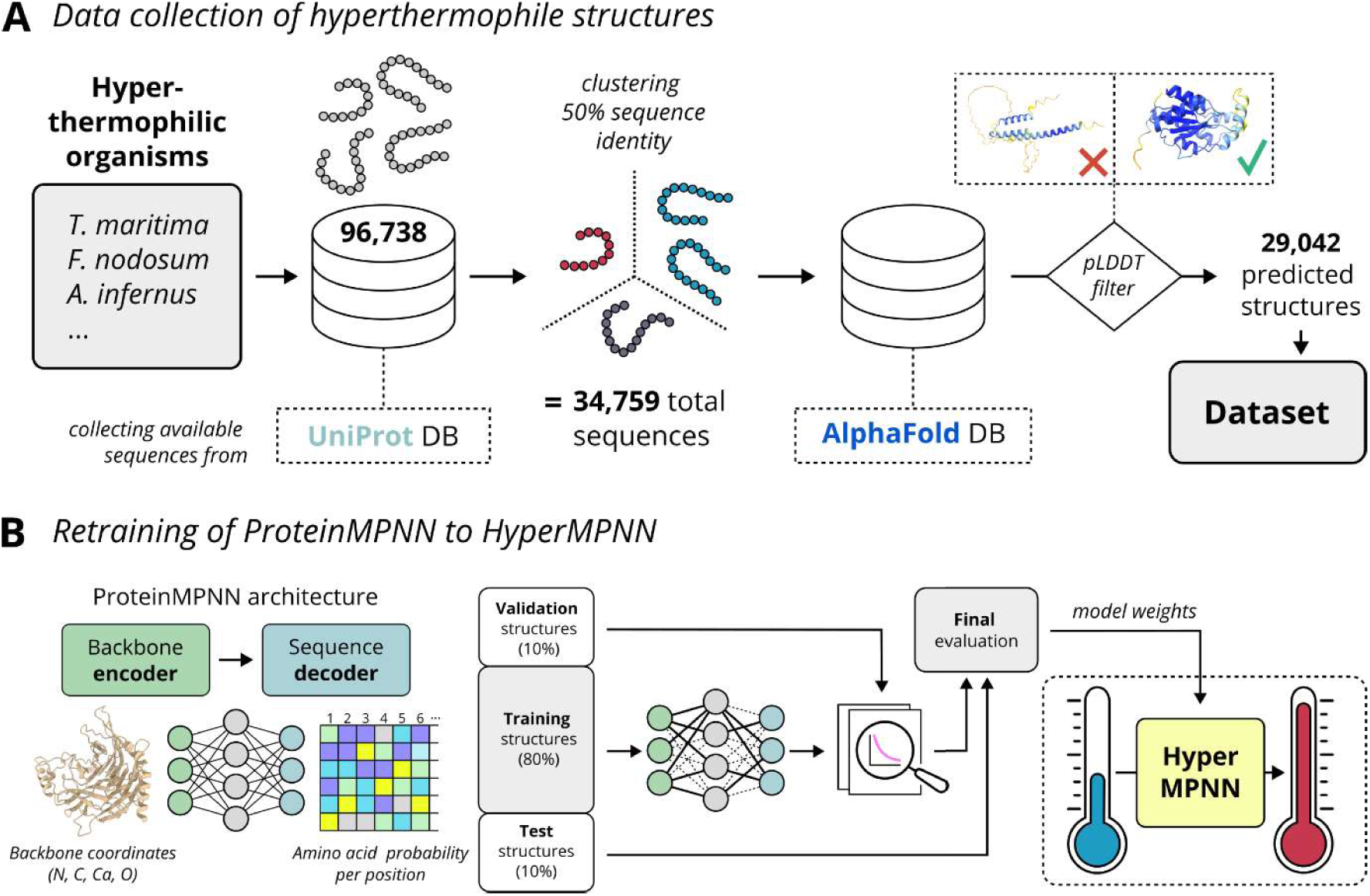
General overview of the approach. **A)** Sequences of hyperthermophiles were collected from the UniProt database and clustered to 50% sequence identity. The AlphaFold database was used to collect predicted structures which were filtered by local and overall pLDDT, resulting in 29,042 predicted structures. **B)** ProteinMPNN was retrained to HyperMPNN using the collected hyperthermophile dataset and then compared to the original model.

Next, the amino acid composition was analyzed, comparing these proteins to proteins from the mesophile *E. coli* (**Fig. 2**). Although a similar analysis has previously been performed on sequence level^10,21^, our focus on predicted structures now allowed us to analyze structural properties of the proteins. It could be observed that the core of proteins from hyperthermophiles has 4.4% more apolar residues than the *E. coli* reference. For the surface, proteins from hyperthermophiles had a 3.9% increase in positively charged residues, a 4.1% increase in apolar residues, a 4.6% reduction in polar residues and a 4.6% reduction in others. For the core, an 4.4% increase in apolar residues in proteins from hyperthermophiles was observed.

**Fig. 2.**
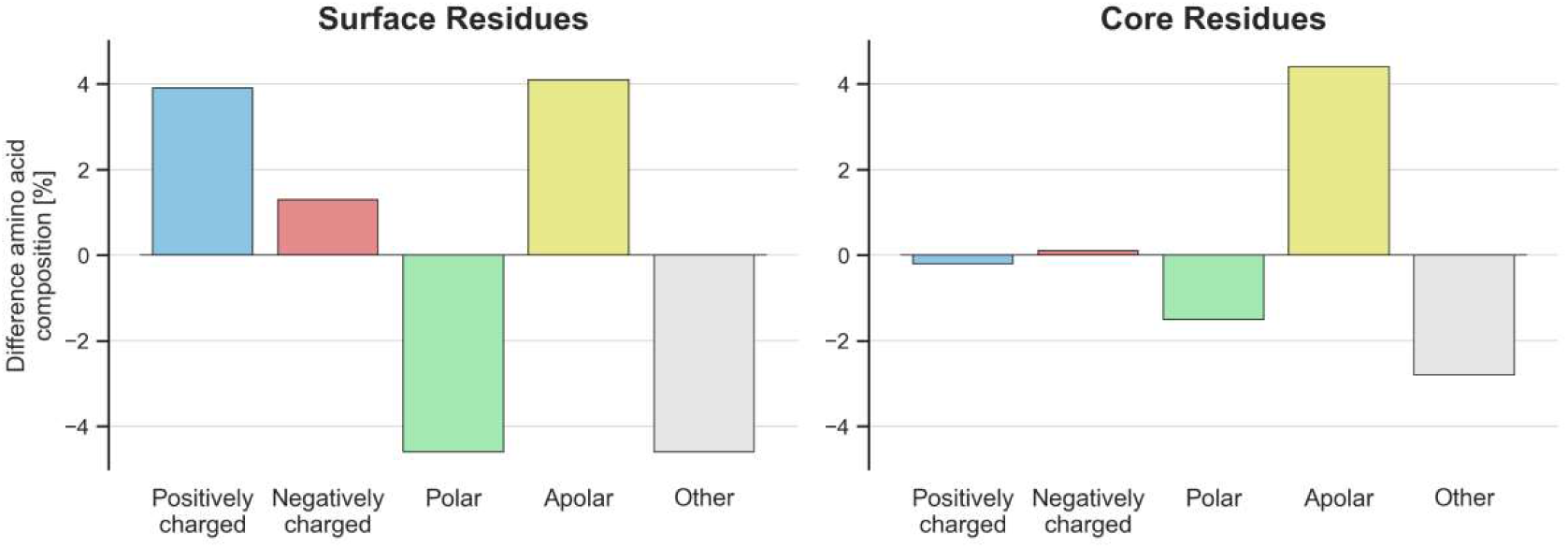
Differences in amino acid composition of proteins from hyperthermophilic and mesophilic organisms. The difference refers to %Hyperthemophilic - %Mesophilic amino acid composition. Protein residues are grouped in surface or core residues depending on whether their solvent-accessible surface area is above 30Å. **Positively charged**: Arg, Lys. **Negatively charged:** Asp, Glu. **Polar:** Asn, Gln, Ser, Thr. **Apolar:** Iso, Leu, Met, Phe, Trp, Tyr, Val. **Others:** Ala, Cys, Gly, His, Pro.

### ProteinMPNN fails to recover the unique amino acid composition of proteins from hyperthermophilies

To probe whether ProteinMPNN is agnostic to the source organism of a protein, the amino acid composition was analyzed after redesign of all proteins from our thermophile dataset (**Fig. 3, top**). Compared to the original surface composition of proteins from hyperthermophiles, there was an increase in alanine, proline, and glutamate, and a decrease in arginine, serine, and isoleucine. Interestingly, redesigning proteins from *E. coli* with ProteinMPNN resulted in almost the identical amino acid composition as designing on proteins from thermophiles, with an even greater bias towards alanine (**Fig. 3, bottom**). In general, the resulting sequences do not follow the original amino acid composition. For *E. coli* there was a large increase in charged lysine and glutamate residues on the surface of resulting proteins, a ProteinMPNN bias observed previously^7^. In comparison, we also used a protein language model (PLM) from the Evolutionary Scale Modeling (ESM)^22^ family to redesign all proteins from thermophiles, with the resulting sequences closely matching the original frequencies (**Fig. S1**).

**Fig. 3.**
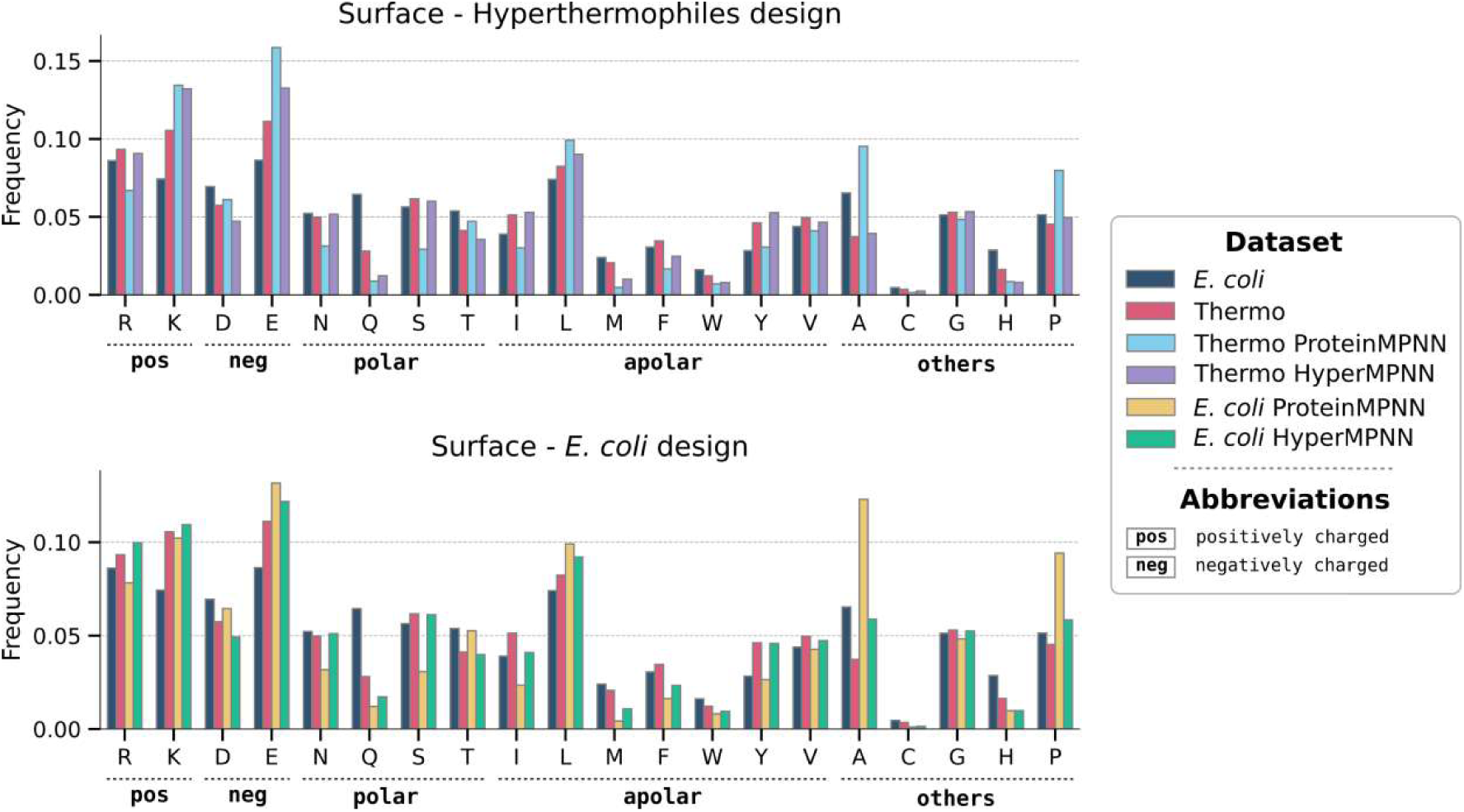
ProteinMPNN fails to recover the unique amino acid composition of hyperthermophiles. Barplots of amino acid frequencies for surface protein residues identified by solvent-accessible surface area (>30Å). Comparison of the frequencies for proteins from *E. coli* (dark blue), proteins from hyperthermophiles (red) . (**top**) Hyper-thermophile proteins redesigned with ProteinMPNN (light blue) and proteins from hyperthermophiles redesigned with HyperMPNN (violet). (**bottom**) *E. coli* proteins either redesigned with ProteinMPNN (yellow) or the re-trained HyperMPNN (green). The amino acid composition of the core area is shown in suppl. fig. S2.

### Retraining ProteinMPNN on hyperthermophilic proteins allows to optimize for stability

As ProteinMPNN failed to recover the unique amino acid composition of hyperthermophiles, it was retrained and the resulting network termed HyperMPNN. Reproducing the original ProteinMPNN training, the structures were clustered and subsequently split randomly into training, validation, and test clusters. During training, a random member from each cluster of the training set was picked and fed into the training loop. The model was trained with different amounts of added Gaussian noise (**Fig. S3**). However, the resulting model perplexity of 5.183 (exponentiated categorical cross-entropy loss per residue) and accuracy of 0.483 (percentage of correct amino acids recovered) for the test clusters were comparable to the original ProteinMPNN results using a model trained with 0.02 backbone noise as reference^15^. The hyperparameter optimization was performed using a grid search approach, but did not result in a substantial improvement compared to the standard model architecture and selected batch sizes, number of epochs etc. (**Table S3**).

After training, HyperMPNN was used to design either on the *E. coli* or on the thermophile protein dataset (**Fig. 3**). Similar to ProteinMPNN the resulting amino acid HyperMPNN closely followed the amino acid composition in proteins from hyperthermophiles, fully recovering the observed compositional biases. This includes both the reduction in polar uncharged residues and the increase in apolar residues found at the protein surface. Next, we were interested whether this learned strategy is generally applicable and not limited to proteins specifically found in hyperthermophiles. Therefore, the charge distribution, number of salt bridges, contact order, and radius of gyration of the initial proteins were analyzed in comparison to the redesigned sequences (**Fig. 4**). Both the original proteins from hyperthermophiles and the HyperMPNN redesigned *E. coli* proteins showed a median positive net charge of 1 and 3 respectively. In contrast, both the original *E. coli* proteins and ProteinMPNN redesigns had a negative median net charge (**Fig. 4A**). A common theory is that the added stability of proteins from hyperthermophiles is based on extended salt bridge networks^10,23^. However, we saw no substantial difference in the number of salt bridges between proteins from *E. coli* (median 16.2) or hyperthermophiles (median 17.0). Intriguingly, for redesigns of E. coli proteins the ProteinMPNN designed sequences only had a median of 8.8 salt bridges per protein compared to the 17.0 median of HyperMPNN (**Fig. 4B**). Lastly, both for the contact order (median 38.6 in *E. coli*, median 36.3 in hyperthermophiles) (**Fig. 4C**) and radius of gyration (median 23.1Å in *E. coli*, 22.1Å in hyperthermophiles) (**Fig. 4D**) there was no strong difference between the two sets of proteins.

**Fig 4.**
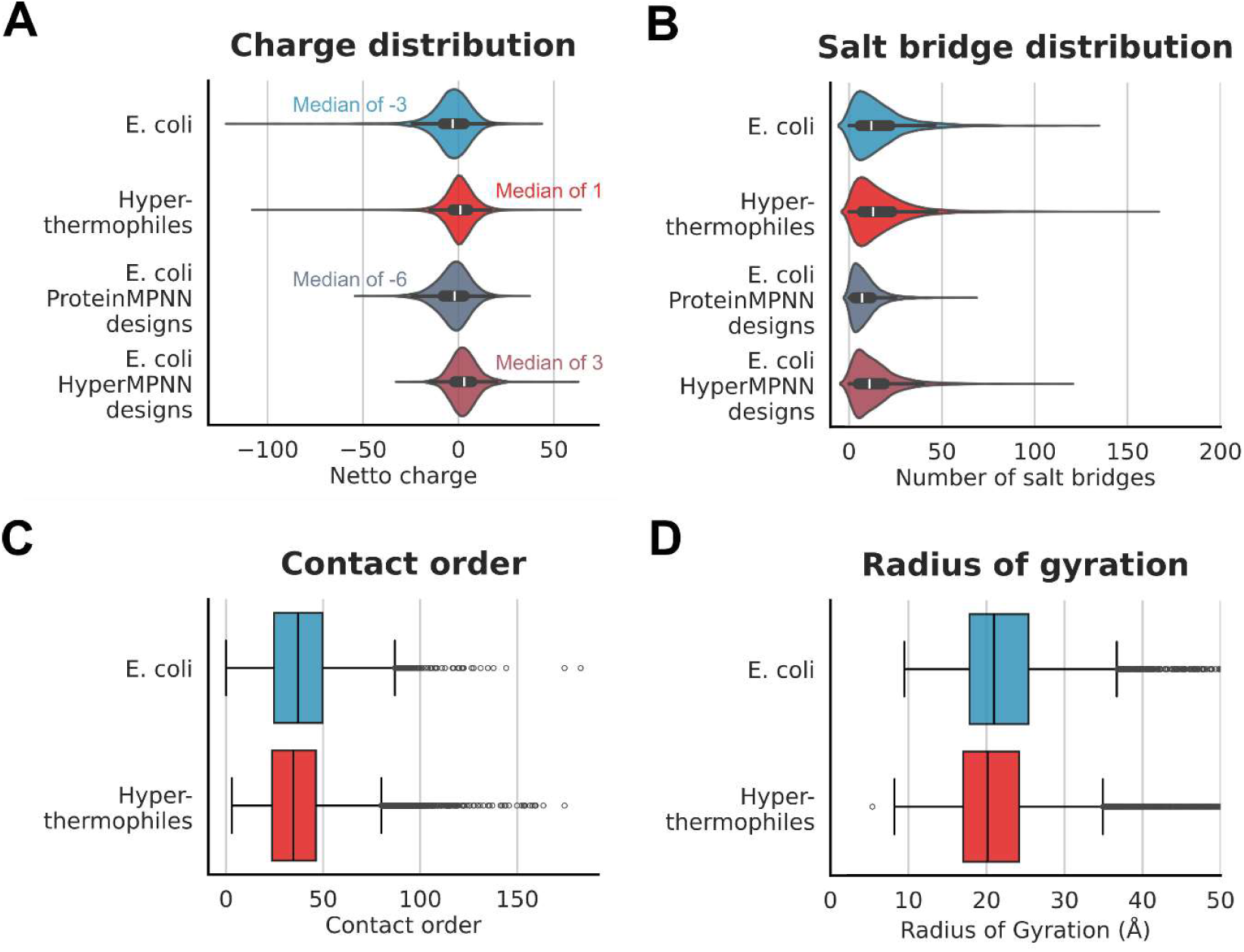
Analysis of proteins from hyperthermophiles and designed proteins. **A)** Difference in charge distribution between proteins from *E. coli*, hyperthermophiles and ProteinMPNN variants. **B)** Comparison of the number of salt bridges in natural and designed proteins. **C)** Overview of contact order in proteins from *E. coli* and thermophiles. **D)** Comparison of the radius of gyration between proteins from *E. coli* and thermophiles. For all plots the median is indicated as a white dot (A) or black line (B, C, D); boxes depict the interquartile range (IQR), the whiskers represent 1.5 x IQR.

### HyperMPNN is an efficient tool to stabilize proteins

To investigate the potential of HyperMPNN in the design of thermostabilized proteins, we employed our design approach to the I53-50B.4PT1 pentamer, which, together with the I53-50A.1PT1 trimer, constitutes a component of the icosahedral I53-50 nanoparticle^14^. Therefore, we redesigned the I53-50B.4PT1 pentamer residues not involved in assembly and used the top 10% of the 200 generated sequences, selected based on their global scores, to derive a consensus sequence (HyperMPNN: I53-50B.HMPNN, ProteinMPNN: I5350B.PMPNN) that was subsequently subjected to experimental validation. The resulting sequence identities to the parent protein were 70% for the HyperMPNN design and 76% for the ProteinMPNN design. Furthermore, the net charge at pH 7 of the designs was calculated in comparison to the parent protein (z_net_ = -0.83), revealing an increase in negative charge for I53-50B.PMPNN (z_net_ = -7.88) and an increase in positive charge for I53-50B.HMPNN (z_net_ = 2.01) (**Fig. 5A**). These findings are further reflected by the surface electrostatic potential (**Fig. S5**), whereas the HyperMPNN design shows an increase of positive charged residues at the lumen-facing regions of the pentamer, while the ProteinMPNN design shows an increase of negative charged residues.

**Fig. 5.**
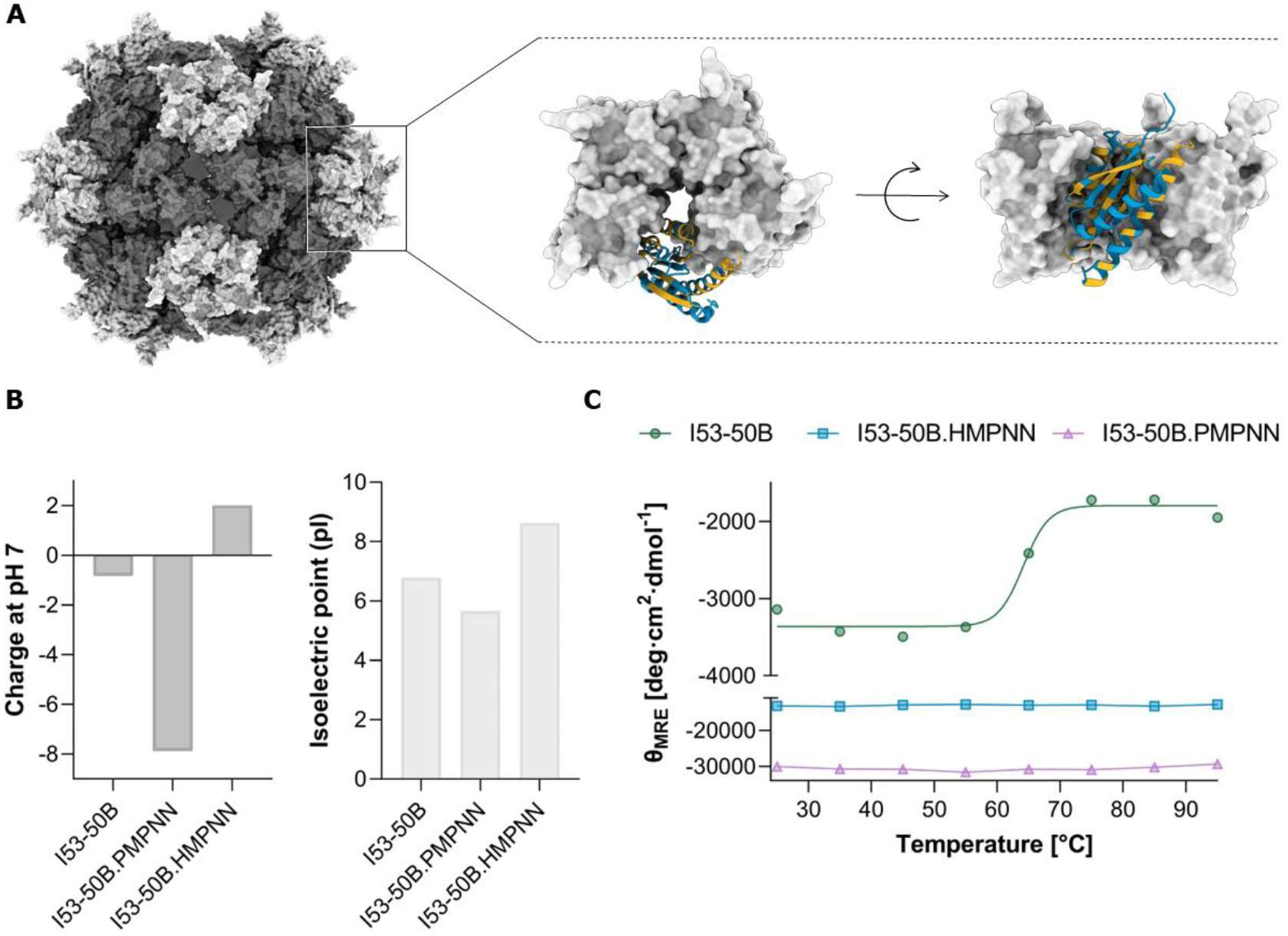
HyperMPNN designs improve thermostability. **A)** I53-50 nanoparticle (PDB: 6P6F^24^) with the zoomed-in view of the I53-50B pentamer. Residues involved in assembly are highlighted in orange, residues set to be designed in blue. **B)** Net charge at pH 7 and isoelectrical point of the parent sequence (I53-50B) and consensus sequences (HyperMPNN: I53-50B.HMPNN, ProteinMPNN: I53-50B.PMPNN). **C** Thermal melting CD plots of the designed and parent I53-50B pentamer measured in mean residue molar ellipticity (Θ_MRE_).

Synthetic DNA carrying the parent gene, the HyperMPNN design, and the ProteinMPNN design were expressed in *E. coli*. The resulting proteins were purified via IMAC followed by SEC. All constructs, including the parent protein, were soluble and eluted as monomers during SEC, whereas the HyperMPNN construct exhibited a lower level of soluble yield in contrast to the parent and ProteinMPNN sequence (I53-50B.HMPNN: 0.4 mg/L culture; I53-50B: 20.4 mg/L culture; I53-50B.PMPNN: 25.8 mg/L culture)

The stability of the proteins was determined using thermal melting circular dichroism (CD). For the parent sequence a melting temperature at 65°C was observed. The HyperMPNN and ProteinMPNN designs remained stable at 95°C (**Fig. 5C** and **S4**), indicating a significant improvement in thermal stability.

## Discussion

In this work, we retrained ProteinMPNN on predicted structures from hyperthermophiles, leveraging their evolutionary adaptation to high temperatures to learn a general strategy for thermal stabilization of proteins (**Fig. 1**). We first compared amino acid frequencies from *E. coli* and hyperthermophiles, finding a bias towards charged residues at the surface and apolar residues in the core of hyperthermophiles (**Fig. 2**). Designing proteins from thermophiles with ProteinMPNN did not recover this unique amino acid composition (**Fig. 3 top**) and therefore we retrained the network on predicted protein structures from hyperthermophiles. In contrast to ProteinMPNN^15^ which was trained on the PDB^25^, HyperMPNN recapitulates the unique amino acid composition of proteins from hyperthermophiles, and successfully transfers it to proteins from other organisms (**Fig. 3 bottom**). As expected, both ProteinMPNN and HyperMPNN are agnostic to the source organism of the input protein and instead apply the same amino acid frequencies derived from their training data. In contrast, as PLMs, such as ESM, are sensitive to the origin of an input sequence their designs remain in the amino acid composition of the source organism (**Fig. S1**). Further, we confirmed that proteins from hyperthermophiles do not have special attributes that would prevent applying the learned strategy to proteins from other organisms (**Fig. 4**), as has been shown previously^10^. Lastly, we experimentally verified our approach by designing the I53-50B.4PT1 pentamer of the icosahedral protein nanoparticle I53-50B.

This study demonstrates that HyperMPNN can substantially enhance the thermal stability of proteins. Specifically, it successfully predicted a variant of I53-50B that remains stable at temperatures up to 95°C, notably surpassing the stability of the parent protein. Given the increasing demand for drug delivery systems capable of withstanding prolonged high temperatures, especially with the rise of new vaccine technologies like mRNA vaccines, our sequence design strategy using HyperMPNN could offer a valuable tool for developing protein nanoparticle variants with enhanced thermal resilience. The lower expression levels of the HyperMPNN construct compared to the parent or ProteinMPNN sequence, may be attributed to several factors, but most probably because of misfolding during synthesis at temperatures that are 30 to 50°C below the natural growth temperature of hyperthermophile organisms^26^. Given the protein’s likely requirement for higher folding temperatures, heterologous expression in *Thermus thermophilus*, a thermophilic host better aligned with the native folding environment, may improve folding, stability, and overall expression yield^27^. From a biotechnological perspective, it would also be interesting to explore HyperMPNN’s potential in designing thermostable enzymes, as high temperature has been identified as a key factor in enhancing the efficiency of industrial operations^28^. Previous studies have shown that mutating the interior surface of I53-50 to positively charged residues facilitates the successful uptake of negatively charged GFP^14^. Therefore, we propose that the I53-50B variant designed by HyperMPNN, which carries an overall positive charge compared to the parent or ProteinMPNN design, may enhance nanoparticle assembly when paired with the I53-50A trimer and negatively charged cargo.

Multiple other studies have focused on the thermal stabilization of proteins, either using biophysical scoring functions^5,6^, evolutionary data^4^ or supervised machine learning approaches^7,8^. While incorporating data from evolutionary sources is very efficient in removing deleterious mutations from the pool of variants^3,4^ and retaining functionality^29^, it requires sufficient data and is only an indirect approach for thermal stabilization. The recent development of high throughput data collection^9^, resulted in higher accuracies for supervised methods directly predicting dG or ddG^7,8^. A relevant example is ThermoMPNN^7^, which uses learned features extracted from ProteinMPNN to directly predict ddG values for mutations given a protein structure. Intriguingly, the authors uncovered a bias in ThermoMPNN predictions favoring the same amino acids we identified as frequently occurring in thermophiles as stabilizing mutations. While this supervised approach is exciting, for now the underlying data has low diversity and is mainly based on small (>100 residues) proteins, limiting generalization to larger folds. In contrast, our sequence dataset from hyperthermophiles spans a wide range of sizes, folds and functionalities as it consists of proteomes from multiple organisms, allowing for the self-supervised learning of a broadly applicable strategy.

The exact mechanisms of adaptation of hyperthermophiles to heat has been an intensively studied subject over time^10,16,17^. Here, a broad dataset of predicted structures from hyperthermophiles is analyzed, going beyond previous single case or sequence-based studies. Similar to previous studies we found a bias towards charged residues at the surface and polar residues in the core of hyperthermophiles^10,21^. Interestingly, we did not find an overall increase in salt bridges, which are commonly associated as driving factors for increased thermal stability^8^.

A limitation of our approach is the use of predicted structures from the AlphaFold database^19,20^. However, the collected predicted structures were filtered by overall and local pLDDT to avoid low quality predictions. Additionally, as our strategy is based on an overall change in the amino acid composition of surface residues, it might require more residues to be redesigned compared to methods focusing on single point mutations. Another potential limitation is the application of HyperMPNN to protein folds which only exist in mammalian organisms, as the underlying training data is from prokaryotes only.

In conclusion, we provide a new tool to (re-)design proteins for high thermal stability, borrowing an evolutionary adoption from hyperthermophilic organisms and in a first proof of concept, we successfully stabilized a vaccine scaffold, improving it’s melting temperature by 30°C.

## Methods

A full protocol capture and code for reproducing the training and results can be found at https://github.com/meilerlab/HyperMPNN.

### Collection of hyperthermophilic protein sequences and predicted structures

Sequences from a list of hyperthermophilic organisms^16,17^ were collected through UniProt^18^. Next, all 96,738 sequences were clustered to 50% sequence identity resulting in 34,759 sequences and their predicted structures were downloaded from the AlphaFold database^19,20^. After filtering for low pLDDT the dataset consisted of 29,042 predicted structures of hyperthermophiles.

### Analysis of amino acid compositions

For the calculation of amino acid frequencies each residue was classified into surface (>30Å) or core (<30Å) as calculated by the Rosetta PerResidueSasaMetric^33^ and its identity counted.

### Retraining of ProteinMPNN and evaluation of HyperMPNN

For retraining ProteinMPNN^15^, the curated dataset of 29,042 predicted single-chain protein structures of hyperthermophiles from the AlphaFold database was used. For every structure, a unique 4-letter pseudo pdb ID and two pytorch data files were created: one containing the general information of the structure (sequence list, date, resolution, etc.) and the second containing chain-specific information (sequence, atom coordinates) (**Table S1**). For the correct parsing of the structures, the `parse_cif_noX.py` was used as a blueprint and a pdb-specific function was applied to extract the needed information for training. The sequences of all predicted structures were clustered again using `mmseqs` with the `easy-cluster` method, 0.3 sequence identity (`--min-seq-id` flag), 0.8 covered residues (`-c` flag), and coverage mode 0 (`--cov-mode` flag) to map representative and member sequences. The identified clusters were randomly split into training (13,426), validation (897), and testing (552) clusters using `train_test_split` from the `sklearn` package. Before training, some functions provided in `utils.py` from the original ProteinMPNN training directory were slightly changed to fit our selected folder structure and some preprocessing steps, but no other changes were made to the training loop from the original implementation^16^. For training, 0.2 Å of Gaussian noise were added to the backbone coordinates, dropout of 10%, and the network was trained for 300 epochs with a batch size of 10,000 and 20,000 examples per epoch. For other parameters, the default values were used (**Table S2**). grid search was performed to find better hyperparameters by varying batch size, examples per epoch, number of nearest neighbors for the sparse graph, number of hidden dimensions, and dropout (**Table S3**) by trying to train the network without running into overfitting and analyzing the achieved accuracy and perplexity.

ProteinMPNN and HyperMPNN designs of proteins from *E. coli* or hyperthermophiles were created by sampling one sequence per protein at temperature of 0.001. The resulting sequences were threaded on the input structures using the Rosetta `SimpleThreadingMover` with five rounds of packing. The amino acid composition of the resulting designs was calculated as described above. Metrics were calculated through RosettaScripts^33,34^ or PyRosetta^35^ using the `NetCharge` filter and the `SaltBridgeCalculator` with default values. For the radius of gyration and contact order the corresponding Rosetta energy terms were used^36^.

### Sequence design of I53-50B.4PT1

For experimental testing of the HyperMPNN model one oligomeric protein component of a self-assembling protein nanoparticle was selected, specifically the pentameric component I53-50B.4PT1 (PDB: 6P6F^24^). The chains B, C, D, E, and F were extracted and renumbered. The target residues for design were selected using Rosetta `InterfaceByVector` mover with a 10 Å CB distance cutoff for the interface between chains B-C and B-F. All residues of chain B that are not located at the interfaces were taken into account for sequence design. Sequences were generated from the crystal structure using HyperMPNN and ProteinMPNN, producing 200 designs per target with a sampling temperature of 0.001. Subsequently, sequences were sorted according to their global_score, and the consensus sequence was derived from the top 10%, by calculating the most common amino acid at each position (see https://github.com/meilerlab/HyperMPNN for the scripts).

### Protein expression and purification

The codon-optimized DNA fragments encoding for the designed and native sequences with C-terminal hexahistidine tags were purchased from Integrated DNA Technologies (IDT) as gBlock^TM^ Gene Fragments. Genes with suitable overhangs for *Nde*I/*Not*I restriction digest were cloned into a pET21a(+) target vector following the Gibson Assembly cloning protocol (NEB # E2611S).

The assembled plasmids were employed for the transformation of *E. coli* BL21(DE3) via heat shock, whereas the DNA was incubated with the cells for 30 minutes on ice, followed by a 45 second heat shock at 42°C and a 2 minute incubation on ice. Afterwards 150 μL Luria-Broth (LB) medium was added and samples were incubated for 1 hour at 37°C and 550 rpm on an Eppendorf ThermoMixer^®^. Subsequently, the cells were plated on agar plates containing 100 μg/mL ampicillin and were incubated overnight at 37°C. Single colonies were selected and used to inoculate 100 μg/mL ampicillin, which was then incubated overnight at 37°C and 210 rpm. Afterwards, 10 mL of the preculture was used to inoculate 1L of LB medium (100 mg/mL for plasmid extraction following the ZymoPURE Miniprep protocol (Zymo #D4210). The purified plasmid was sent to Sanger sequencing (Azenta/GeneWiz) for sequence verification. The 1 L cultures were grown at 37°C and 210 rpm until an OD_600_ of 0.6-0.8 was reached. Protein production was induced by the addition of 0.5 mM IPTG, and the cultures were incubated for a 4 hour period at 37°C with agitation at 210 rpm. Cells were harvested by centrifugation at 10,628 x g for 20 minutes and resuspended in lysis buffer (50 mM Tris-HCl, 150 mM NaCl, 10 mM imidazole, 0.1 mg/mL lysozyme, pH 8.0). Cell disruption was achieved through sonication (2 min, 10s pulse, 10s off, 60% amplitude). The lysate was separated from cell debris by centrifugation at 42,511 x g for 30 minutes and subsequently applied to a Ni-NTA resin (Qiagen) in a gravity flow column, which was equilibrated with lysis buffer. The hexahistidine-tagged proteins were allowed to bind for 30 minutes at room temperature, after which the resin was washed with 8 column volumes (CV) wash buffer (50 mM Tris-HCl, 150 mM NaCl, 20 mM imidazole, pH 8.0). Proteins were eluted with 3 CV elution buffer (50 mM Tris-HCl, 150 mM NaCl, 250 mM imidazole, pH 8.0) and subsequently subjected to size exclusion chromatography (SEC) using a HiLoad Superdex 16/600 200 pg column (GE Healthcare) on an ÄKTA pure^TM^(GE Healthcare) at a flow rate of 1.0 mL/min. Fractions containing the protein of interest were analyzed using SDS-PAGE, pooled concentrators (Satorius). Concentration was determined by measuring the absorbance at 280 nm using a spectrophotometer (NanoDrop^TM^ One). Purified proteins were stored in TBS buffer(50 mM Tris-HCl, 150 mM NaCl, pH 8.0) at -20°C until subjected to further analysis.

### Measuring protein melting temperature

The thermostability of the protein designs were assessed using circular dichroism (CD) measurements performed on a JASCO J-1500 machine with a 1 mm pathlength cuvette. Purified proteins were prepared at a concentration of 5 µM in TBS buffer, and the sample temperatures were increased from 25°C to 95°C. Full spectrum scans were recorded at 10°C intervals, spanning the range from 190 nm to 260 nm. The normalized signal at 223 nm was plotted against temperature and fitted to a Boltzman sigmoidal curve using GraphPad Prism 9. Melting points were determined from inflection points of the resulting curves.

## Availability and resources

Code for reproducing the results can be found at https://github.com/meilerlab/HyperMPNN.

## Acknowledgment

Computations for this work were done (in part) using resources of the Leipzig University Computing Centre. We would like to thank the Google DeepMind team for sharing their protein structure predictions. Additionally, we would like to thank the ProteinMPNN authors for making their tool available to the community. Moreover, we wish to thank Prof. Dr. Anette Beck-Sickinger (Institute of Biochemistry, Leipzig University) for allowing us to conduct measurements at their CD spectrometer.

## Author contributions

Conceptualization: ME JM CTS. Data Curation: ME MB PS. Formal Analysis: ME PS MB. Funding Acquisition: CTS LK JM. Investigation: ME MB PS. Methodology: ME PS MB CT JM. Project Administration: ME CTS. Resources: CTS LK JM. Software: ME MB. Supervision: CTS LK JM. Validation: ME MB PS. Visualization: ME PS MB. Writing - Original Draft Preparation: ME PS MB. Writing - Review & Editing: ME MB PS CTS LK JM.

