## Supplemental Information for "HyperMPNN – A general strategy to design thermostable proteins learned from hyperthermophiles"

### Supplementary Information

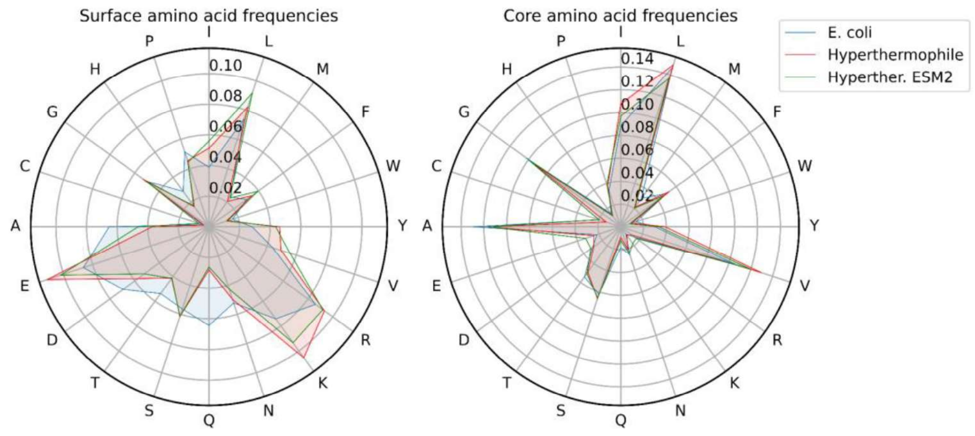


**Fig. S1. Radar plot of amino acid frequencies for protein residues grouped by solvent-accessible surface area in core (<30Å) or surface (>30Å).** Comparison of the frequencies for proteins from *E. coli* (blue), hyperthermophiles (red), or hyperthermophilic proteins redesigned with ESM (green).

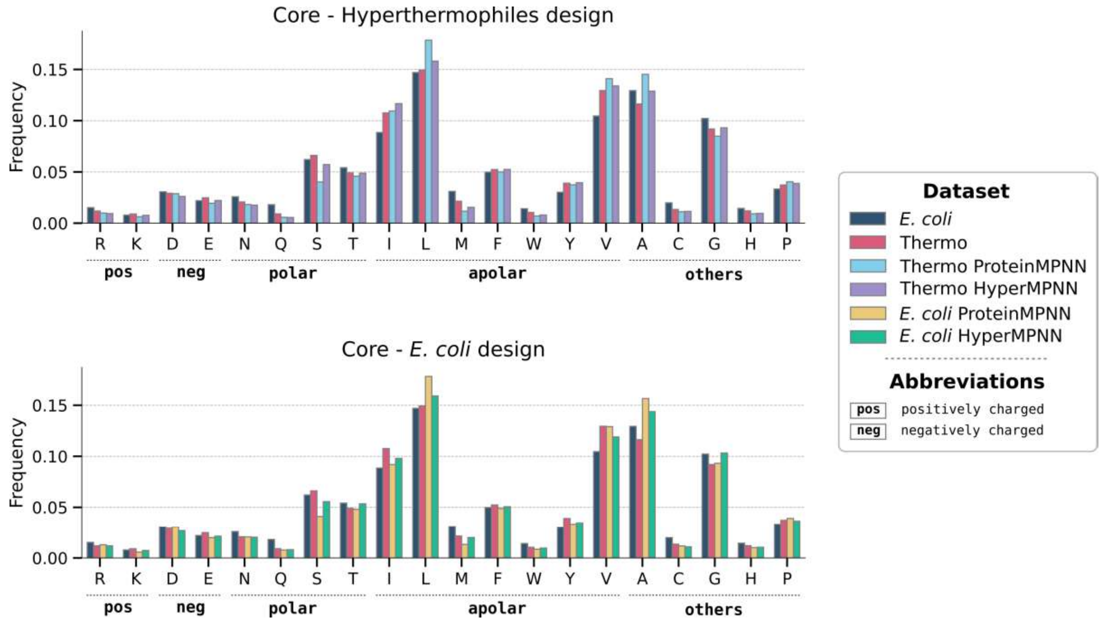


**Fig. 3. ProteinMPNN fails to recover the unique amino acid composition of hyperthermophiles.** Barplots of amino acid frequencies for core protein residues identified by solvent-accessible surface area (<30Å). Comparison of the frequencies for proteins from *E. coli* (dark blue), proteins from hyperthermophiles (red) . (**top**) Hyper- thermophile proteins redesigned with ProteinMPNN (light blue) and proteins from hyperthermophiles redesigned with HyperMPNN (violet). (**bottom**) *E. coli* proteins either redesigned with ProteinMPNN (yellow) or the re-trained HyperMPNN (green).

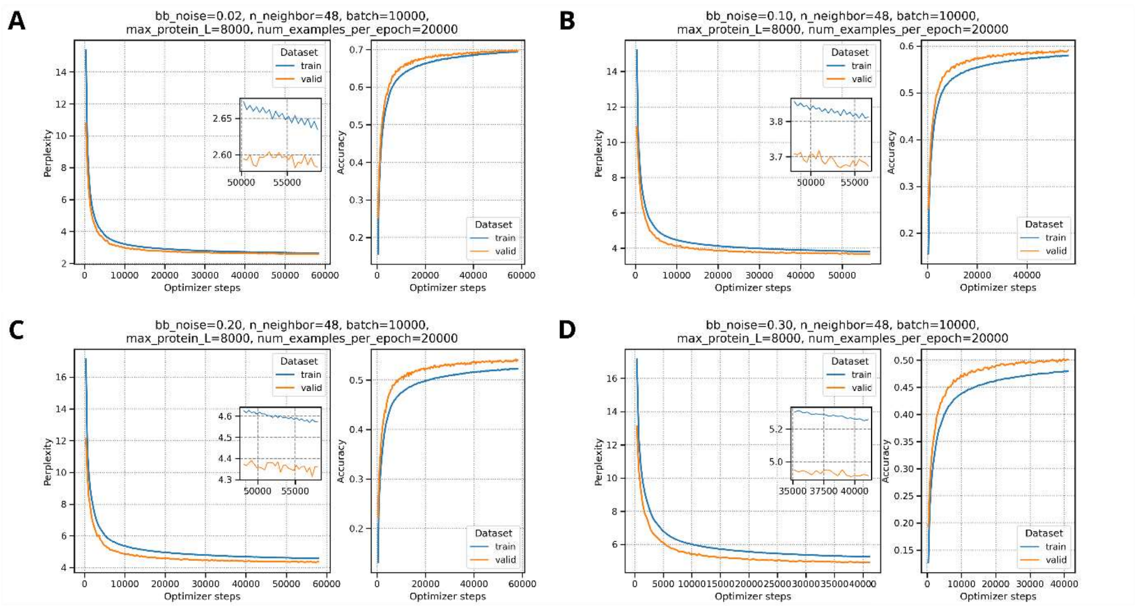


**Fig. S3. Training results for HyperMPNN with different values of added backbone noise.** Other parameters were held fixed for the training runs. A full list of parameters is shown in Table S2. The amount of backbone noise added during training is for (A) 0.02, (B) 0.10, (C) 0.20, and (D) 0.30.

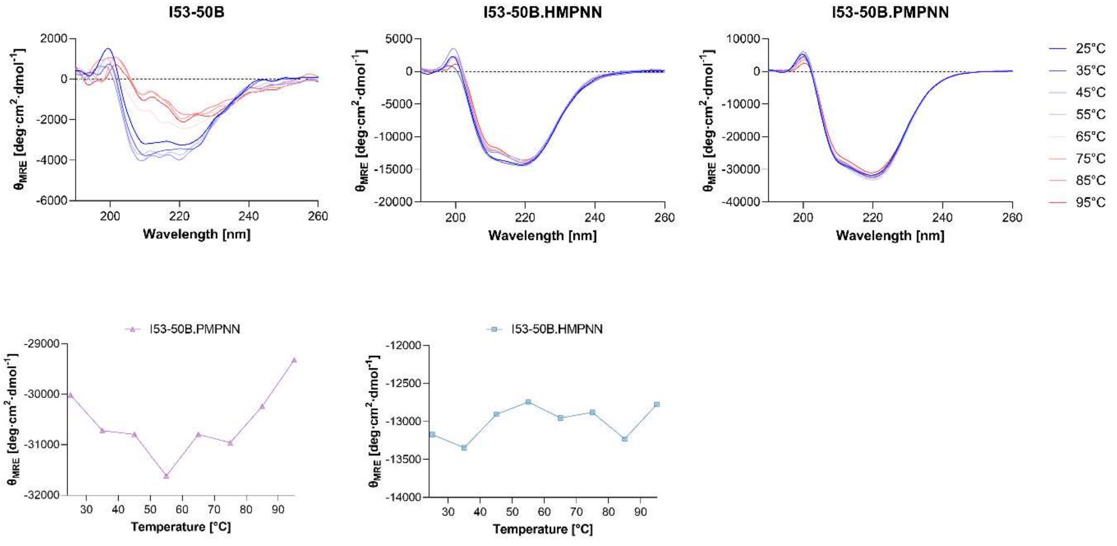


**Fig. S4. Thermal melting CD spectra of I53-50B designs and parent sequence.** Spectra were recorded over a temperature gradient from 25°C to 95°C. The CD signal is reported as mean residue molar ellipticity $\Theta_{MRE}$. The bottom two diagrams illustrate the normalized signal at 223 nm of the two designs plotted against temperature, indicating no detectable thermal transition. I53-50B.hypCS represent the HyperMPNN and I5350B.protCS the ProteinMPNN designed (consensus) sequence.

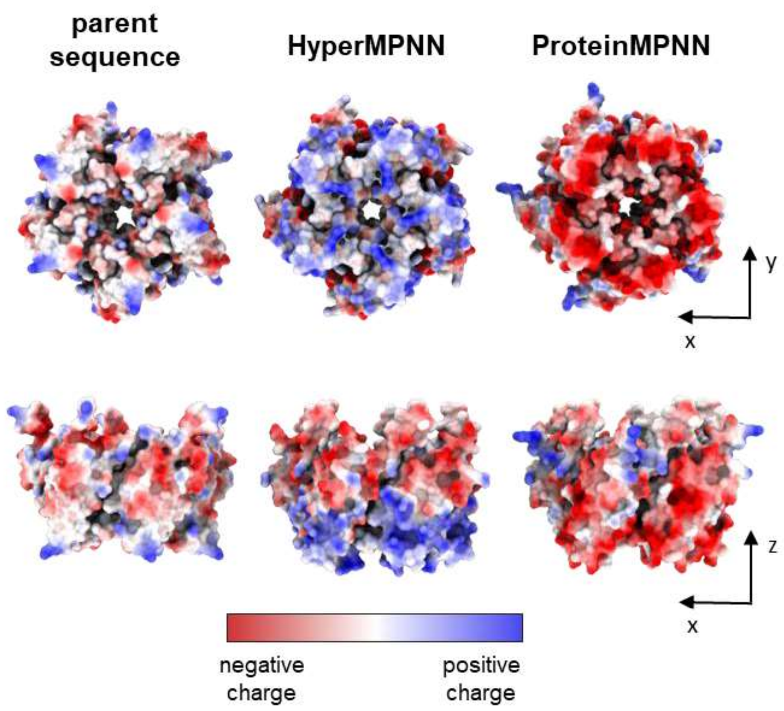


**Fig. S5. Surface charge distribution of I53-50B and designs.** The protein surface is color-coded according to the electrostatic potential, from negative charges (red) to positive charges (blue). The determination of charges was conducted using the Amber 20 built-in of ChimeraX.

**Tab. S1. Extracted information from the pdb input structures which are mandatory for the training script to function correctly.** The information is stored in pytorch-specific `pt` files (two data files per structure). Pseudo values, mandatory fields that are not used here during training, have a default value. The field with the comment containing `'pdbx_struct_assembly_gen'` is also not used because we only use single-chain pdbs.

| Data files | Field | Datatype | Default | Comment |
|---|---|---|---|---|
| **General structure file** | seq | `list[list[str]]` | - | List of sequences for the whole structure. |
| | method | `str` | `'biopython'` | NMR, etc. |
| | date | `str` | `'2023-01-01'` | Pseudo creation date |
| | resolution | `float` | `0.0` | - |
| | chains | `list[str]` | `['A']` | All chains in structure. |
| | id | `str` | - | Pseudo four letter id created. |
| | asmb_chains | `list[str]` | `['A']` | CIF field, `pdbx_struct_assembly_gen.asym_id_list` |
| | asmb_details | `list[str]` | `['own_parsing']` | CIF field, `pdbx_struct_assembly_gen.details` |
| | asmb_method | `list[str]` | `['none']` | CIF field, `pdbx_struct_assembly_gen.method_details` |

| | | | | |
|---|---|---|---|---|
| | asmd_ids | `list[str]` | `['1']` | CIF field, `pdbx_struct_assembly_gen.assembly_id` |
| | asmb_xform0 | `tensor` | $I_4$, Identity matrix | - |
| | tm | `tensor` | `[[[1., 1., 0]]]` | Output of TMalign |
| **Chain specific file** | seq | `str` | - | Sequence of chain/structure |
| | xyz | `tensor` Shape: `Lx14x3` | - | XYZ coordinates of every atom in every sidechain, max. lengths 2. dim = 14 (Tryptophan) |

**Tab S2. The setting used for the training script provided in the ProteinMPNN GitHub to train HyperMPNN with structures from hyperthermophilic proteins and hyper parameter settings.** Values that differ from the default settings are marked with an asterisk (*). If the parameter was changed during the hyper parameter search, the values will be listed in the last column.

| Flag | Value | Hyperparameter settings |
|---|---|---|
| `num_epochs` | 300 (*) | - |
| `save_model_every_n_epochs` | 10 | - |
| `reload_data_every_n_epochs` | 2 | - |
| `num_examples_per_epoch` | 1000 (*) | [15000, 20000, 35000] |
| `batch_size` | 2000 (*) | [8000, 10000, 15000] |
| `max_protein_length` | 5000 (*) | - |
| `hidden_dim` | 128 | [96, 128, 156] |
| `num_encoder_layers` | 3 | - |
| `num_decoder_layers` | 3 | - |
| `num_neighbors` | 48 | [38, 48, 58] |
| `dropout` | 0.1 | [0.05, 0.1, 0.15] |
| `backbone_noise` | 0.2 | [0.1] |
| `rescut` | 3.5 | - |
| `gradient_norm` | -1.0 | - |
| `mixed_precision` | True | - |
| `debug` | False | - |

**Tab S3. Results for the hyper parameter grid search.** The values for perplexity train/valid, overfitting (bool), and the accuracy for the training and validation set correspond to the final evaluation at the end of the training loop.

| batch size | Examples per epoch | Number of neighbors | Hidden Dim. | Dropout | Total time steps | perplexity train | perplexity valid | Over-fitting | accuracy train | accuracy valid |
|---|---|---|---|---|---|---|---|---|---|---|
| 8000 | 15000 | 38 | 96 | 0.05 | 87386 | 3.907 | 3.768 | FALSE | 0.572 | 0.583 |
| 8000 | 15000 | 38 | 96 | 0.1 | 87394 | 4.008 | 3.764 | FALSE | 0.564 | 0.581 |
| 8000 | 15000 | 38 | 96 | 0.15 | 87390 | 4.083 | 3.763 | FALSE | 0.558 | 0.581 |
| 8000 | 15000 | 38 | 128 | 0.05 | 87390 | 3.666 | 3.734 | TRUE | 0.592 | 0.585 |
| 8000 | 15000 | 38 | 128 | 0.1 | 87392 | 3.768 | 3.668 | FALSE | 0.583 | 0.589 |
| 8000 | 15000 | 38 | 128 | 0.15 | 87388 | 3.846 | 3.665 | FALSE | 0.577 | 0.589 |
| 8000 | 15000 | 38 | 156 | 0.05 | 87392 | 3.481 | 3.764 | TRUE | 0.609 | 0.583 |
| 8000 | 15000 | 38 | 156 | 0.1 | 87394 | 3.601 | 3.647 | TRUE | 0.598 | 0.592 |
| 8000 | 15000 | 38 | 156 | 0.15 | 87396 | 3.692 | 3.607 | FALSE | 0.59 | 0.594 |
| 8000 | 15000 | 48 | 96 | 0.05 | 87390 | 3.901 | 3.76 | FALSE | 0.572 | 0.582 |
| 8000 | 15000 | 48 | 96 | 0.1 | 87390 | 4 | 3.762 | FALSE | 0.564 | 0.581 |
| 8000 | 15000 | 48 | 96 | 0.15 | 87394 | 4.072 | 3.786 | FALSE | 0.559 | 0.58 |
| 8000 | 15000 | 48 | 128 | 0.05 | 87390 | 3.658 | 3.712 | TRUE | 0.593 | 0.587 |
| 8000 | 15000 | 48 | 128 | 0.1 | 87394 | 3.761 | 3.688 | FALSE | 0.584 | 0.589 |
| 8000 | 15000 | 48 | 128 | 0.15 | 87394 | 3.849 | 3.658 | FALSE | 0.577 | 0.591 |
| 8000 | 15000 | 48 | 156 | 0.05 | 87396 | 3.478 | 3.737 | TRUE | 0.61 | 0.585 |
| 8000 | 15000 | 48 | 156 | 0.1 | 87386 | 3.59 | 3.636 | TRUE | 0.599 | 0.593 |
| 8000 | 15000 | 48 | 156 | 0.15 | 87394 | 3.687 | 3.595 | FALSE | 0.59 | 0.596 |
| 8000 | 15000 | 58 | 96 | 0.05 | 87392 | 3.907 | 3.753 | FALSE | 0.572 | 0.583 |
| 8000 | 15000 | 58 | 96 | 0.1 | 87392 | 3.994 | 3.74 | FALSE | 0.565 | 0.584 |
| 8000 | 15000 | 58 | 96 | 0.15 | 87376 | 4.085 | 3.76 | FALSE | 0.558 | 0.582 |
| 8000 | 15000 | 58 | 128 | 0.05 | 87396 | 3.653 | 3.709 | TRUE | 0.593 | 0.587 |

| | | | | | | | | | | |
|---|---|---|---|---|---|---|---|---|---|---|
| 8000 | 15000 | 58 | 128 | 0.1 | 87394 | 3.769 | 3.657 | FALSE | 0.583 | 0.591 |
| 8000 | 15000 | 58 | 128 | 0.15 | 87392 | 3.845 | 3.646 | FALSE | 0.577 | 0.591 |
| 8000 | 15000 | 58 | 156 | 0.05 | 87382 | 3.473 | 3.724 | TRUE | 0.61 | 0.587 |
| 8000 | 15000 | 58 | 156 | 0.1 | 87390 | 3.601 | 3.656 | TRUE | 0.598 | 0.59 |
| 8000 | 15000 | 58 | 156 | 0.15 | 87386 | 3.691 | 3.586 | FALSE | 0.59 | 0.596 |
| 8000 | 20000 | 38 | 96 | 0.05 | 87392 | 3.909 | 3.768 | FALSE | 0.571 | 0.581 |
| 8000 | 20000 | 38 | 96 | 0.1 | 87392 | 4.006 | 3.772 | FALSE | 0.564 | 0.58 |
| 8000 | 20000 | 38 | 96 | 0.15 | 87394 | 4.078 | 3.749 | FALSE | 0.558 | 0.581 |
| 8000 | 20000 | 38 | 128 | 0.05 | 87396 | 3.666 | 3.706 | TRUE | 0.592 | 0.588 |
| 8000 | 20000 | 38 | 128 | 0.1 | 87394 | 3.769 | 3.652 | FALSE | 0.583 | 0.59 |
| 8000 | 20000 | 38 | 128 | 0.15 | 87384 | 3.845 | 3.654 | FALSE | 0.577 | 0.59 |
| 8000 | 20000 | 38 | 156 | 0.05 | 87394 | 3.483 | 3.75 | TRUE | 0.609 | 0.586 |
| 8000 | 20000 | 38 | 156 | 0.1 | 87394 | 3.598 | 3.628 | TRUE | 0.599 | 0.593 |
| 8000 | 20000 | 38 | 156 | 0.15 | 87398 | 3.688 | 3.608 | FALSE | 0.59 | 0.594 |
| 8000 | 20000 | 48 | 96 | 0.05 | 87390 | 3.904 | 3.76 | FALSE | 0.572 | 0.582 |
| 8000 | 20000 | 48 | 96 | 0.1 | 87384 | 3.995 | 3.73 | FALSE | 0.565 | 0.584 |
| 8000 | 20000 | 48 | 96 | 0.15 | 87392 | 4.079 | 3.737 | FALSE | 0.558 | 0.583 |
| 8000 | 20000 | 48 | 128 | 0.05 | 87390 | 3.65 | 3.711 | TRUE | 0.594 | 0.587 |
| 8000 | 20000 | 48 | 128 | 0.1 | 87392 | 3.759 | 3.644 | FALSE | 0.584 | 0.591 |
| 8000 | 20000 | 48 | 128 | 0.15 | 87394 | 3.845 | 3.645 | FALSE | 0.577 | 0.593 |
| 8000 | 20000 | 48 | 156 | 0.05 | 87390 | 3.475 | 3.749 | TRUE | 0.61 | 0.585 |
| 8000 | 20000 | 48 | 156 | 0.1 | 87388 | 3.598 | 3.648 | TRUE | 0.598 | 0.592 |
| 8000 | 20000 | 48 | 156 | 0.15 | 87400 | 3.69 | 3.59 | FALSE | 0.59 | 0.595 |
| 8000 | 20000 | 58 | 96 | 0.05 | 87394 | 3.909 | 3.749 | FALSE | 0.572 | 0.584 |
| 8000 | 20000 | 58 | 96 | 0.1 | 87386 | 4.003 | 3.761 | FALSE | 0.564 | 0.583 |
| 8000 | 20000 | 58 | 96 | 0.15 | 87392 | 4.074 | 3.745 | FALSE | 0.559 | 0.583 |
| 8000 | 20000 | 58 | 128 | 0.05 | 87388 | 3.653 | 3.709 | TRUE | 0.593 | 0.588 |
| 8000 | 20000 | 58 | 128 | 0.15 | 87384 | 3.845 | 3.641 | FALSE | 0.577 | 0.593 |
| 8000 | 20000 | 58 | 156 | 0.05 | 87394 | 3.477 | 3.727 | TRUE | 0.609 | 0.587 |
| 8000 | 20000 | 58 | 156 | 0.1 | 87394 | 3.587 | 3.628 | TRUE | 0.599 | 0.593 |
| 8000 | 20000 | 58 | 156 | 0.15 | 87382 | 3.687 | 3.592 | FALSE | 0.591 | 0.596 |
| 8000 | 35000 | 38 | 96 | 0.05 | 87388 | 3.917 | 3.776 | FALSE | 0.571 | 0.581 |
| 8000 | 35000 | 38 | 96 | 0.1 | 87388 | 4.008 | 3.754 | FALSE | 0.564 | 0.582 |
| 8000 | 35000 | 38 | 96 | 0.15 | 87396 | 4.085 | 3.764 | FALSE | 0.558 | 0.581 |
| 8000 | 35000 | 38 | 128 | 0.05 | 87390 | 3.666 | 3.718 | TRUE | 0.592 | 0.586 |
| 8000 | 35000 | 38 | 128 | 0.1 | 87392 | 3.771 | 3.653 | FALSE | 0.583 | 0.591 |
| 8000 | 35000 | 38 | 128 | 0.15 | 87390 | 3.853 | 3.645 | FALSE | 0.576 | 0.592 |
| 8000 | 35000 | 38 | 156 | 0.05 | 87388 | 3.483 | 3.724 | TRUE | 0.609 | 0.587 |
| 8000 | 35000 | 38 | 156 | 0.1 | 87394 | 3.592 | 3.662 | TRUE | 0.599 | 0.59 |
| 8000 | 35000 | 38 | 156 | 0.15 | 87388 | 3.687 | 3.605 | FALSE | 0.59 | 0.595 |
| 8000 | 35000 | 48 | 96 | 0.05 | 87394 | 3.899 | 3.747 | FALSE | 0.573 | 0.585 |
| 8000 | 35000 | 48 | 96 | 0.1 | 87394 | 3.999 | 3.732 | FALSE | 0.564 | 0.585 |
| 8000 | 35000 | 48 | 96 | 0.15 | 87392 | 4.081 | 3.763 | FALSE | 0.558 | 0.582 |
| 8000 | 35000 | 48 | 128 | 0.05 | 87390 | 3.65 | 3.691 | TRUE | 0.593 | 0.589 |
| 8000 | 35000 | 48 | 128 | 0.1 | 87390 | 3.767 | 3.661 | FALSE | 0.583 | 0.592 |
| 8000 | 35000 | 48 | 128 | 0.15 | 87398 | 3.843 | 3.64 | FALSE | 0.577 | 0.592 |
| 8000 | 35000 | 48 | 156 | 0.05 | 87392 | 3.483 | 3.731 | TRUE | 0.609 | 0.586 |
| 8000 | 35000 | 48 | 156 | 0.1 | 87384 | 3.597 | 3.628 | TRUE | 0.598 | 0.592 |
| 8000 | 35000 | 48 | 156 | 0.15 | 87398 | 3.685 | 3.594 | FALSE | 0.59 | 0.597 |
| 8000 | 35000 | 58 | 96 | 0.05 | 87390 | 3.895 | 3.733 | FALSE | 0.573 | 0.584 |
| 8000 | 35000 | 58 | 96 | 0.1 | 87392 | 3.991 | 3.724 | FALSE | 0.565 | 0.586 |
| 8000 | 35000 | 58 | 96 | 0.15 | 87392 | 4.086 | 3.734 | FALSE | 0.558 | 0.585 |
| 8000 | 35000 | 58 | 128 | 0.05 | 87392 | 3.649 | 3.689 | TRUE | 0.594 | 0.59 |
| 8000 | 35000 | 58 | 128 | 0.1 | 87390 | 3.76 | 3.649 | FALSE | 0.584 | 0.591 |
| 8000 | 35000 | 58 | 128 | 0.15 | 87396 | 3.847 | 3.643 | FALSE | 0.577 | 0.591 |
| 8000 | 35000 | 58 | 156 | 0.05 | 87396 | 3.475 | 3.724 | TRUE | 0.61 | 0.587 |
| 8000 | 35000 | 58 | 156 | 0.1 | 87390 | 3.584 | 3.635 | TRUE | 0.6 | 0.593 |
| 8000 | 35000 | 58 | 156 | 0.15 | 87394 | 3.685 | 3.595 | FALSE | 0.591 | 0.596 |
| 10000 | 15000 | 38 | 96 | 0.05 | 70266 | 3.92 | 3.785 | FALSE | 0.571 | 0.58 |
| 10000 | 15000 | 38 | 96 | 0.1 | 70266 | 4.011 | 3.763 | FALSE | 0.563 | 0.582 |
| 10000 | 15000 | 38 | 96 | 0.15 | 70290 | 4.091 | 3.763 | FALSE | 0.557 | 0.581 |
| 10000 | 15000 | 38 | 128 | 0.05 | 70276 | 3.666 | 3.726 | TRUE | 0.592 | 0.587 |
| 10000 | 15000 | 38 | 128 | 0.1 | 70256 | 3.766 | 3.67 | FALSE | 0.583 | 0.59 |
| 10000 | 15000 | 38 | 128 | 0.15 | 70260 | 3.839 | 3.642 | FALSE | 0.577 | 0.592 |
| 10000 | 15000 | 38 | 156 | 0.05 | 70264 | 3.47 | 3.751 | TRUE | 0.61 | 0.585 |
| 10000 | 15000 | 38 | 156 | 0.1 | 70250 | 3.593 | 3.664 | TRUE | 0.599 | 0.59 |
| 10000 | 15000 | 38 | 156 | 0.15 | 70260 | 3.682 | 3.605 | FALSE | 0.591 | 0.594 |
| 10000 | 15000 | 48 | 96 | 0.05 | 70266 | 3.903 | 3.775 | FALSE | 0.572 | 0.581 |
| 10000 | 15000 | 48 | 96 | 0.1 | 70272 | 4.005 | 3.762 | FALSE | 0.564 | 0.582 |
| 10000 | 15000 | 48 | 96 | 0.15 | 70266 | 4.085 | 3.75 | FALSE | 0.558 | 0.584 |
| 10000 | 15000 | 48 | 128 | 0.05 | 70284 | 3.653 | 3.713 | TRUE | 0.594 | 0.587 |
| 10000 | 15000 | 48 | 128 | 0.1 | 70276 | 3.757 | 3.662 | FALSE | 0.585 | 0.591 |
| 10000 | 15000 | 48 | 128 | 0.15 | 70258 | 3.835 | 3.63 | FALSE | 0.578 | 0.592 |
| 10000 | 15000 | 48 | 156 | 0.05 | 70266 | 3.458 | 3.75 | TRUE | 0.611 | 0.585 |
| 10000 | 15000 | 48 | 156 | 0.1 | 70258 | 3.59 | 3.656 | TRUE | 0.599 | 0.592 |
| 10000 | 15000 | 48 | 156 | 0.15 | 70264 | 3.679 | 3.609 | FALSE | 0.591 | 0.595 |
| 10000 | 15000 | 58 | 96 | 0.05 | 70282 | 3.899 | 3.783 | FALSE | 0.573 | 0.581 |
| 10000 | 15000 | 58 | 96 | 0.1 | 70260 | 3.996 | 3.746 | FALSE | 0.565 | 0.584 |
| 10000 | 15000 | 58 | 96 | 0.15 | 70264 | 4.079 | 3.755 | FALSE | 0.558 | 0.583 |
| 10000 | 15000 | 58 | 128 | 0.05 | 70260 | 3.646 | 3.719 | TRUE | 0.594 | 0.587 |
| 10000 | 15000 | 58 | 128 | 0.1 | 70274 | 3.76 | 3.659 | FALSE | 0.584 | 0.592 |
| 10000 | 15000 | 58 | 128 | 0.15 | 70260 | 3.843 | 3.635 | FALSE | 0.577 | 0.593 |

| | | | | | | | | | | |
|---|---|---|---|---|---|---|---|---|---|---|
| 10000 | 15000 | 58 | 156 | 0.05 | 70278 | 3.47 | 3.73 | TRUE | 0.61 | 0.586 |
| 10000 | 15000 | 58 | 156 | 0.1 | 70264 | 3.587 | 3.66 | TRUE | 0.599 | 0.59 |
| 10000 | 15000 | 58 | 156 | 0.15 | 70256 | 3.679 | 3.586 | FALSE | 0.591 | 0.597 |
| 10000 | 20000 | 38 | 96 | 0.05 | 70284 | 3.913 | 3.761 | FALSE | 0.572 | 0.583 |
| 10000 | 20000 | 38 | 96 | 0.1 | 70260 | 4.008 | 3.746 | FALSE | 0.564 | 0.584 |
| 10000 | 20000 | 38 | 96 | 0.15 | 70280 | 4.102 | 3.759 | FALSE | 0.557 | 0.583 |
| 10000 | 20000 | 38 | 128 | 0.05 | 70274 | 3.652 | 3.728 | TRUE | 0.594 | 0.585 |
| 10000 | 20000 | 38 | 128 | 0.1 | 70280 | 3.76 | 3.663 | FALSE | 0.584 | 0.59 |
| 10000 | 20000 | 38 | 128 | 0.15 | 70264 | 3.85 | 3.652 | FALSE | 0.577 | 0.591 |
| 10000 | 20000 | 38 | 156 | 0.05 | 70256 | 3.469 | 3.745 | TRUE | 0.61 | 0.585 |
| 10000 | 20000 | 38 | 156 | 0.1 | 70262 | 3.592 | 3.666 | TRUE | 0.599 | 0.59 |
| 10000 | 20000 | 38 | 156 | 0.15 | 70268 | 3.688 | 3.62 | FALSE | 0.59 | 0.594 |
| 10000 | 20000 | 48 | 96 | 0.05 | 70266 | 3.903 | 3.756 | FALSE | 0.572 | 0.584 |
| 10000 | 20000 | 48 | 96 | 0.1 | 70266 | 3.994 | 3.739 | FALSE | 0.565 | 0.584 |
| 10000 | 20000 | 48 | 96 | 0.15 | 70272 | 4.078 | 3.761 | FALSE | 0.559 | 0.584 |
| 10000 | 20000 | 48 | 128 | 0.05 | 70270 | 3.663 | 3.724 | TRUE | 0.592 | 0.586 |
| 10000 | 20000 | 48 | 128 | 0.1 | 70262 | 3.759 | 3.658 | FALSE | 0.584 | 0.591 |
| 10000 | 20000 | 48 | 128 | 0.15 | 70266 | 3.846 | 3.632 | FALSE | 0.577 | 0.593 |
| 10000 | 20000 | 48 | 156 | 0.05 | 70270 | 3.458 | 3.745 | TRUE | 0.612 | 0.586 |
| 10000 | 20000 | 48 | 156 | 0.1 | 70266 | 3.588 | 3.655 | TRUE | 0.599 | 0.593 |
| 10000 | 20000 | 48 | 156 | 0.15 | 70278 | 3.682 | 3.614 | FALSE | 0.591 | 0.595 |
| 10000 | 20000 | 58 | 96 | 0.05 | 70252 | 3.901 | 3.751 | FALSE | 0.572 | 0.583 |
| 10000 | 20000 | 58 | 96 | 0.1 | 70278 | 4.005 | 3.738 | FALSE | 0.564 | 0.585 |
| 10000 | 20000 | 58 | 96 | 0.15 | 70262 | 4.085 | 3.737 | FALSE | 0.558 | 0.583 |
| 10000 | 20000 | 58 | 128 | 0.05 | 70272 | 3.656 | 3.706 | TRUE | 0.593 | 0.588 |
| 10000 | 20000 | 58 | 128 | 0.1 | 70288 | 3.752 | 3.656 | FALSE | 0.585 | 0.591 |
| 10000 | 20000 | 58 | 128 | 0.15 | 70268 | 3.841 | 3.617 | FALSE | 0.577 | 0.593 |
| 10000 | 20000 | 58 | 156 | 0.05 | 70260 | 3.46 | 3.733 | TRUE | 0.611 | 0.587 |
| 10000 | 20000 | 58 | 156 | 0.1 | 70284 | 3.579 | 3.67 | TRUE | 0.6 | 0.59 |
| 10000 | 20000 | 58 | 156 | 0.15 | 70274 | 3.679 | 3.604 | FALSE | 0.591 | 0.595 |
| 10000 | 35000 | 38 | 96 | 0.05 | 70264 | 3.913 | 3.775 | FALSE | 0.572 | 0.583 |
| 10000 | 35000 | 38 | 96 | 0.1 | 70274 | 4.007 | 3.754 | FALSE | 0.564 | 0.583 |
| 10000 | 35000 | 38 | 96 | 0.15 | 70280 | 4.087 | 3.768 | FALSE | 0.558 | 0.581 |
| 10000 | 35000 | 38 | 128 | 0.05 | 70278 | 3.664 | 3.725 | TRUE | 0.593 | 0.587 |
| 10000 | 35000 | 38 | 128 | 0.1 | 70266 | 3.759 | 3.669 | FALSE | 0.584 | 0.589 |
| 10000 | 35000 | 38 | 128 | 0.15 | 70270 | 3.847 | 3.662 | FALSE | 0.577 | 0.589 |
| 10000 | 35000 | 38 | 156 | 0.05 | 70284 | 3.47 | 3.757 | TRUE | 0.61 | 0.585 |
| 10000 | 35000 | 38 | 156 | 0.1 | 70258 | 3.598 | 3.663 | TRUE | 0.598 | 0.59 |
| 10000 | 35000 | 38 | 156 | 0.15 | 70262 | 3.683 | 3.61 | FALSE | 0.591 | 0.596 |
| 10000 | 35000 | 48 | 96 | 0.05 | 70258 | 3.906 | 3.76 | FALSE | 0.572 | 0.584 |
| 10000 | 35000 | 48 | 96 | 0.1 | 70290 | 3.998 | 3.721 | FALSE | 0.564 | 0.586 |
| 10000 | 35000 | 48 | 96 | 0.15 | 70252 | 4.092 | 3.77 | FALSE | 0.557 | 0.582 |
| 10000 | 35000 | 48 | 128 | 0.05 | 70270 | 3.652 | 3.713 | TRUE | 0.594 | 0.588 |
| 10000 | 35000 | 48 | 128 | 0.1 | 70284 | 3.757 | 3.644 | FALSE | 0.584 | 0.591 |
| 10000 | 35000 | 48 | 128 | 0.15 | 70288 | 3.839 | 3.643 | FALSE | 0.578 | 0.592 |
| 10000 | 35000 | 48 | 156 | 0.05 | 70276 | 3.466 | 3.749 | TRUE | 0.611 | 0.586 |
| 10000 | 35000 | 48 | 156 | 0.1 | 70278 | 3.584 | 3.645 | TRUE | 0.599 | 0.592 |
| 10000 | 35000 | 48 | 156 | 0.15 | 70264 | 3.681 | 3.603 | FALSE | 0.591 | 0.596 |
| 10000 | 35000 | 58 | 96 | 0.05 | 70290 | 3.908 | 3.77 | FALSE | 0.572 | 0.583 |
| 10000 | 35000 | 58 | 96 | 0.1 | 70268 | 4.004 | 3.745 | FALSE | 0.564 | 0.582 |
| 10000 | 35000 | 58 | 96 | 0.15 | 70270 | 4.086 | 3.776 | FALSE | 0.558 | 0.581 |
| 10000 | 35000 | 58 | 128 | 0.05 | 70282 | 3.657 | 3.714 | TRUE | 0.593 | 0.587 |
| 10000 | 35000 | 58 | 128 | 0.1 | 70284 | 3.756 | 3.658 | FALSE | 0.584 | 0.592 |
| 10000 | 35000 | 58 | 128 | 0.15 | 70274 | 3.836 | 3.632 | FALSE | 0.578 | 0.593 |
| 10000 | 35000 | 58 | 156 | 0.05 | 70272 | 3.47 | 3.74 | TRUE | 0.61 | 0.586 |
| 10000 | 35000 | 58 | 156 | 0.1 | 70288 | 3.581 | 3.654 | TRUE | 0.6 | 0.591 |
| 10000 | 35000 | 58 | 156 | 0.15 | 70274 | 3.678 | 3.611 | FALSE | 0.591 | 0.594 |
| 15000 | 15000 | 38 | 96 | 0.05 | 47302 | 3.935 | 3.806 | FALSE | 0.569 | 0.579 |
| 15000 | 15000 | 38 | 96 | 0.1 | 47316 | 4.027 | 3.768 | FALSE | 0.562 | 0.582 |
| 15000 | 15000 | 38 | 96 | 0.15 | 47312 | 4.113 | 3.773 | FALSE | 0.556 | 0.58 |
| 15000 | 15000 | 38 | 128 | 0.05 | 47318 | 3.67 | 3.746 | TRUE | 0.592 | 0.585 |
| 15000 | 15000 | 38 | 128 | 0.1 | 47322 | 3.776 | 3.669 | FALSE | 0.583 | 0.59 |
| 15000 | 15000 | 38 | 128 | 0.15 | 47296 | 3.861 | 3.664 | FALSE | 0.576 | 0.59 |
| 15000 | 15000 | 38 | 156 | 0.05 | 47302 | 3.474 | 3.816 | TRUE | 0.61 | 0.58 |
| 15000 | 15000 | 38 | 156 | 0.1 | 47296 | 3.589 | 3.696 | TRUE | 0.599 | 0.588 |
| 15000 | 15000 | 38 | 156 | 0.15 | 47302 | 3.691 | 3.626 | FALSE | 0.59 | 0.594 |
| 15000 | 15000 | 48 | 96 | 0.05 | 47304 | 3.924 | 3.785 | FALSE | 0.571 | 0.581 |
| 15000 | 15000 | 48 | 96 | 0.1 | 47308 | 4.01 | 3.739 | FALSE | 0.563 | 0.584 |
| 15000 | 15000 | 48 | 96 | 0.15 | 47318 | 4.086 | 3.755 | FALSE | 0.558 | 0.582 |
| 15000 | 15000 | 48 | 128 | 0.05 | 47318 | 3.663 | 3.736 | TRUE | 0.593 | 0.585 |
| 15000 | 15000 | 48 | 128 | 0.1 | 47308 | 3.766 | 3.669 | FALSE | 0.583 | 0.59 |
| 15000 | 15000 | 48 | 128 | 0.15 | 47302 | 3.839 | 3.63 | FALSE | 0.578 | 0.592 |
| 15000 | 15000 | 48 | 156 | 0.05 | 47300 | 3.464 | 3.814 | TRUE | 0.611 | 0.578 |
| 15000 | 15000 | 48 | 156 | 0.1 | 47300 | 3.588 | 3.66 | TRUE | 0.599 | 0.591 |
| 15000 | 15000 | 48 | 156 | 0.15 | 47314 | 3.681 | 3.613 | FALSE | 0.591 | 0.594 |
| 15000 | 15000 | 58 | 96 | 0.05 | 47302 | 3.918 | 3.763 | FALSE | 0.571 | 0.583 |
| 15000 | 15000 | 58 | 96 | 0.1 | 47292 | 4.012 | 3.777 | FALSE | 0.564 | 0.583 |
| 15000 | 15000 | 58 | 96 | 0.15 | 47316 | 4.095 | 3.752 | FALSE | 0.557 | 0.582 |
| 15000 | 15000 | 58 | 128 | 0.05 | 47312 | 3.656 | 3.727 | TRUE | 0.593 | 0.586 |
| 15000 | 15000 | 58 | 128 | 0.1 | 47306 | 3.768 | 3.681 | FALSE | 0.583 | 0.589 |
| 15000 | 15000 | 58 | 128 | 0.15 | 47302 | 3.845 | 3.66 | FALSE | 0.577 | 0.59 |
| 15000 | 15000 | 58 | 156 | 0.15 | 47306 | 3.677 | 3.603 | FALSE | 0.592 | 0.595 |

| | | | | | | | | | | |
|---|---|---|---|---|---|---|---|---|---|---|
| 15000 | 20000 | 38 | 96 | 0.05 | 47298 | 3.921 | 3.787 | FALSE | 0.571 | 0.581 |
| 15000 | 20000 | 38 | 96 | 0.1 | 47308 | 4.032 | 3.762 | FALSE | 0.562 | 0.582 |
| 15000 | 20000 | 38 | 96 | 0.15 | 47300 | 4.106 | 3.763 | FALSE | 0.556 | 0.58 |
| 15000 | 20000 | 38 | 128 | 0.05 | 47292 | 3.668 | 3.737 | TRUE | 0.592 | 0.585 |
| 15000 | 20000 | 38 | 128 | 0.1 | 47306 | 3.775 | 3.679 | FALSE | 0.583 | 0.589 |
| 15000 | 20000 | 38 | 128 | 0.15 | 47280 | 3.855 | 3.652 | FALSE | 0.576 | 0.591 |
| 15000 | 20000 | 38 | 156 | 0.05 | 47316 | 3.463 | 3.769 | TRUE | 0.611 | 0.582 |
| 15000 | 20000 | 38 | 156 | 0.1 | 47294 | 3.598 | 3.705 | TRUE | 0.599 | 0.587 |
| 15000 | 20000 | 38 | 156 | 0.15 | 47296 | 3.681 | 3.635 | FALSE | 0.591 | 0.591 |
| 15000 | 20000 | 48 | 96 | 0.05 | 47318 | 3.922 | 3.812 | FALSE | 0.571 | 0.58 |
| 15000 | 20000 | 48 | 96 | 0.1 | 47310 | 4.018 | 3.772 | FALSE | 0.563 | 0.583 |
| 15000 | 20000 | 48 | 96 | 0.15 | 47284 | 4.093 | 3.762 | FALSE | 0.557 | 0.581 |
| 15000 | 20000 | 48 | 128 | 0.05 | 47286 | 3.67 | 3.749 | TRUE | 0.592 | 0.583 |
| 15000 | 20000 | 48 | 128 | 0.1 | 47312 | 3.773 | 3.682 | FALSE | 0.583 | 0.588 |
| 15000 | 20000 | 48 | 128 | 0.15 | 47306 | 3.851 | 3.652 | FALSE | 0.576 | 0.592 |
| 15000 | 20000 | 48 | 156 | 0.05 | 47308 | 3.461 | 3.788 | TRUE | 0.611 | 0.581 |
| 15000 | 20000 | 48 | 156 | 0.1 | 47292 | 3.577 | 3.672 | TRUE | 0.6 | 0.589 |
| 15000 | 20000 | 48 | 156 | 0.15 | 47298 | 3.674 | 3.609 | FALSE | 0.591 | 0.594 |
| 15000 | 20000 | 58 | 96 | 0.05 | 47304 | 3.918 | 3.766 | FALSE | 0.571 | 0.583 |
| 15000 | 20000 | 58 | 96 | 0.1 | 47312 | 4.012 | 3.747 | FALSE | 0.563 | 0.584 |
| 15000 | 20000 | 58 | 96 | 0.15 | 47310 | 4.099 | 3.752 | FALSE | 0.557 | 0.583 |
| 15000 | 20000 | 58 | 128 | 0.05 | 47306 | 3.656 | 3.758 | TRUE | 0.593 | 0.583 |
| 15000 | 20000 | 58 | 128 | 0.1 | 47312 | 3.768 | 3.669 | FALSE | 0.584 | 0.59 |
| 15000 | 20000 | 58 | 128 | 0.15 | 47302 | 3.846 | 3.659 | FALSE | 0.577 | 0.589 |
| 15000 | 20000 | 58 | 156 | 0.05 | 47288 | 3.464 | 3.784 | TRUE | 0.611 | 0.582 |
| 15000 | 35000 | 38 | 96 | 0.05 | 47322 | 3.931 | 3.799 | FALSE | 0.57 | 0.581 |
| 15000 | 35000 | 38 | 96 | 0.1 | 47312 | 4.018 | 3.769 | FALSE | 0.563 | 0.582 |
| 15000 | 35000 | 38 | 96 | 0.15 | 47302 | 4.104 | 3.791 | FALSE | 0.556 | 0.579 |
| 15000 | 35000 | 38 | 128 | 0.05 | 47294 | 3.673 | 3.744 | TRUE | 0.592 | 0.584 |
| 15000 | 35000 | 38 | 128 | 0.1 | 47304 | 3.773 | 3.701 | FALSE | 0.583 | 0.587 |
| 15000 | 35000 | 38 | 128 | 0.15 | 47314 | 3.858 | 3.671 | FALSE | 0.576 | 0.588 |
| 15000 | 35000 | 38 | 156 | 0.05 | 47292 | 3.47 | 3.772 | TRUE | 0.61 | 0.584 |
| 15000 | 35000 | 38 | 156 | 0.1 | 47304 | 3.6 | 3.686 | TRUE | 0.598 | 0.589 |
| 15000 | 35000 | 38 | 156 | 0.15 | 47298 | 3.689 | 3.613 | FALSE | 0.59 | 0.594 |
| 15000 | 35000 | 48 | 96 | 0.05 | 47292 | 3.923 | 3.785 | FALSE | 0.57 | 0.581 |
| 15000 | 35000 | 48 | 96 | 0.1 | 47310 | 4.011 | 3.741 | FALSE | 0.564 | 0.583 |
| 15000 | 35000 | 48 | 96 | 0.15 | 47292 | 4.097 | 3.743 | FALSE | 0.557 | 0.584 |
| 15000 | 35000 | 48 | 128 | 0.05 | 47282 | 3.659 | 3.751 | TRUE | 0.593 | 0.584 |
| 15000 | 35000 | 48 | 128 | 0.1 | 47296 | 3.766 | 3.665 | FALSE | 0.584 | 0.591 |
| 15000 | 35000 | 48 | 128 | 0.15 | 47304 | 3.845 | 3.649 | FALSE | 0.577 | 0.591 |
| 15000 | 35000 | 48 | 156 | 0.05 | 47314 | 3.469 | 3.785 | TRUE | 0.611 | 0.583 |
| 15000 | 35000 | 48 | 156 | 0.1 | 47304 | 3.588 | 3.661 | TRUE | 0.599 | 0.592 |
| 15000 | 35000 | 48 | 156 | 0.15 | 47302 | 3.685 | 3.631 | FALSE | 0.59 | 0.592 |
| 15000 | 35000 | 58 | 96 | 0.05 | 47296 | 3.914 | 3.789 | FALSE | 0.571 | 0.581 |
| 15000 | 35000 | 58 | 96 | 0.1 | 47302 | 4.011 | 3.765 | FALSE | 0.563 | 0.582 |
| 15000 | 35000 | 58 | 96 | 0.15 | 47310 | 4.094 | 3.742 | FALSE | 0.557 | 0.583 |
| 15000 | 35000 | 58 | 128 | 0.05 | 47308 | 3.658 | 3.74 | TRUE | 0.593 | 0.586 |
| 15000 | 35000 | 58 | 128 | 0.1 | 47298 | 3.763 | 3.699 | FALSE | 0.584 | 0.588 |
| 15000 | 35000 | 58 | 128 | 0.15 | 47302 | 3.847 | 3.637 | FALSE | 0.577 | 0.592 |
| 15000 | 35000 | 58 | 156 | 0.15 | 32636 | 3.772 | 3.66 | FALSE | 0.583 | 0.59 |

584
